# A calibrated novelty flag for fungal ITS metabarcoding: choosing the error rate at which sequences are declared new

**DOI:** 10.64898/2026.07.29.741524

**Authors:** Aaron O’Brien, César Marín, Pilar Parada

## Abstract

1. Environmental fungal surveys routinely recover internal transcribed spacer (ITS) sequences that cannot be assigned at fine taxonomic ranks, the so-called fungal “dark matter.” Such sequences are set aside by thresholding a similarity or confidence score at a conventional value. Those conventions do abstain, but the error rate a threshold implies is neither stated nor selectable, and a threshold defined for one kind of score does not transfer to another.
2. We present a conformal novelty flag that supplies what is missing: each query receives a *p*-value with a distribution-free guarantee that the rate of falsely declaring a known sequence novel is bounded by a user-chosen *α*. We evaluate it on a leave-one-genus-out benchmark built from UNITE, on alignment identity, a *k*-mer bootstrap consensus and two neural classifiers’ output probabilities, and on a soil fungal dataset.
3. The flag holds its nominal rate across two orders of magnitude in *α*, so an operating point can be chosen rather than inherited: at *α* = 0.05 it fires on 4.3% of known-genus queries and recovers 19.2% of genuinely novel genera. The cutoff holding a 5% error rate here is 64.8% identity, nowhere near the customary 97%, showing how little a threshold carries its error rate between datasets. Coverage transferred across eleven settings spanning those four scores, two amplicon regions and a sevenfold change in reference size, all within 1.1 percentage points of nominal, while detection ranged from 5.2% to 53.3%: the guarantee is on the error rate and not on power, and two of our settings are valid but uninformative. Applied to soil data the flag identifies 20.7% of amplicon sequence variants as novel at a controlled 5% error rate, 16.3% under an abundance filter. Half of those recur near-identically among GlobalFungi’s unnamed environmental variants while fewer than one in ten matches a named species hypothesis, a sixfold skew towards the uncatalogued against 1.9-fold for sequences the flag passes.
4. The flag turns an arbitrary cutoff into a decision with a stated error rate, and in doing so converts dark matter from a residue into a set of prioritizable targets for formal description.

## 1 Introduction

Fungi are central to terrestrial ecosystem function, yet our census of fungal diversity remains radically incomplete. Estimates of global fungal richness range from 2.2–3.8 million species [10] to as many as 6.28 million [4], against roughly 155,000 that have been formally described [22]. Metabarcoding of the ITS region routinely recovers sequences with no close match in curated references such as UNITE [16], environmental “dark matter” novel at the genus level or below, and in many surveys these are the majority of what is recovered; among ectomycorrhizal fungi the unnamed fraction has been put as high as 83% [22]. The practical question is not whether such sequences can be named, since often they cannot, but whether anything reliable can be said about them. Sequence-based frameworks already proceed on that footing: UNITE species hypotheses [11] and the virtual taxa of MaarjAM [18] give unnamed lineages stable identifiers without waiting for names, and thirty fungal lineages have now been formally described from environmental samples and DNA [21]. At minimum one wants to *know* that a sequence is novel, with stated confidence, and to have the analysis decline to name it rather than guess.

Current practice does decline, but informally. A similarity threshold, most often 97% identity though 98% is also in use [15], or a bootstrap confidence cutoff, conventionally 0.8, separates sequences that receive a name from those that do not. That the conventional value is not itself quite settled is already telling: neither figure comes with the error rate it implies. Such conventions are not arbitrary in their consequences, and we show below that a widely used bootstrap cutoff behaves stably across datasets. What they do not provide is a rate. A practitioner who wishes to declare sequences novel while wrongly discarding no more than one known sequence in a hundred has no threshold to reach for, cannot report the rate their chosen threshold actually achieved, and must re-derive the mapping from threshold to rate for every new score they wish to use. The problem is calibration rather than the absence of abstention.

This matters more as scoring methods proliferate. Alignment identity, *k*-mer bootstrap consensus and neural classifier probabilities are all in use and all differ in scale, distribution and meaning, so a threshold established for one carries no interpretation for another. A companion study [17] examines how these scores compare as taxonomic assignment methods and reports that neural classifiers, in particular, all but lose their novelty signal on the ITS2 amplicon that environmental surveys predominantly generate. The present paper is concerned with the complementary question of what to do with whatever score is available.

We therefore adopt inductive conformal anomaly detection [24, 2], which converts any score into a *p*-value with a distribution-free bound on the false-novelty rate, and evaluate it on a leave-one-genus-out benchmark constructed so that the novel class consists of genera genuinely absent from the reference. We use alignment identity as a transparent baseline throughout, verify that the guarantee holds for three further scores of entirely different provenance, and then apply the flag to real soil data to ask what the sequences it flags actually are.

## 2 Related work

Three lines of work bear directly on the problem addressed here, and it is worth being precise about what each already provides.

### Data-derived similarity cutoffs

The inadequacy of a single static threshold is well established, and dnabarcoder [25] addresses it directly by predicting clade-specific cutoffs from a reference dataset rather than adopting a convention, showing that cutoffs vary substantially across the fungal tree and that local cutoffs assign fewer sequences than traditional thresholds while improving accuracy and precision. Each predicted cutoff carries a confidence measure, but that measure quantifies the *resolving power* of the barcode within a clade, estimated by clustering agreement against the reference taxonomy, and the cutoff is selected to maximize it.

It is therefore a property of the clade and the marker, not a per-query error rate, and it is not a quantity a user can set. The distinction matters in a second respect the authors themselves note: their cutoffs are computed from BLAST alignments and would require recomputation for a different search algorithm, whereas the calibration described here applies unchanged to any score. Our contribution is complementary rather than competing: dnabarcoder answers “what cutoff best resolves this clade?” and we answer “what cutoff holds my chosen false-novelty rate, whatever score I am using?”

### Confidence thresholds in classifiers

Established taxonomic classifiers already abstain. RDP-style bootstrap confidence and SINTAX [8] both report a per-query score and withhold assignments beneath a threshold, conventionally 0.8. We show below that this convention behaves stably, so the practice is not defective; what it lacks is a stated and selectable rate, and a threshold meaningful for a bootstrap consensus has no interpretation for an alignment identity or a neural softmax.

### Novelty-aware taxonomic models

A separate strand builds novelty into the model rather than wrapping it afterwards. BayesANT [26] uses Bayesian nonparametric priors so that unobserved taxa may be discovered at each rank, and deep hierarchical Bayesian approaches have been applied to the same problem in insect identification [3]. These quantify uncertainty about placement within a probabilistic model, which is a stronger and more informative output than ours where the model is well specified, and a weaker one where it is not: our guarantee is distribution-free and requires no model of the sequence-generating process, but it also yields only a novelty decision rather than a posterior over taxa. In the same vein, Fujisawa and Imai [9] benchmark out-of-distribution detection for insect barcoding under incomplete references and report that detecting unknown species is harder than identifying known ones, consistent with the moderate separability we find for fungal ITS. Their study and ours differ in marker, kingdom and purpose, but their conclusion that similarity-cutoff-based detection deserves scrutiny is the empirical premise this paper starts from.

Methodologically we build on inductive conformal anomaly detection [24] and on the treatment of conformal *p*-values for outlier testing [5], which establishes the validity of split-conformal *p*-values for exactly this one-sided novelty problem. Our contribution is not to the statistics but to establishing that the guarantee survives contact with real fungal barcode data, four heterogeneous scores and two amplicon regions.

## 3 Implementation

### 3.1 Reference data and ITS2 harmonization

We used the UNITE general release sh_general_release_dynamic_19.02.2025 as reference [1], reduced to the ITS2 subregion, a pool of 99,808 sequences from the release’s 102,137 records, comprising 20,295 reference and 81,842 representative sequences. The general rather than the developer release: the two carry identical headers and so define the same taxonomy, but their sequences differ and only the general release reproduces the ITS2 boundaries recorded for this pool. Because ITS2 is length-variable and its amplicon boundary is primer-dependent, comparing query and reference identity requires that both occupy the same locus. HMM-anchored full-cistron extraction [6] failed on short primer-bounded amplicons, which lack the flanking SSU and 5.8S structure the anchoring model requires. The reference records are full-length and retain that structure, so the reference and the benchmark queries drawn from it were delimited with ITSx [6] directly, giving the 99,808-sequence ITS2 pool from 102,137 input records. The soil ASVs and the GlobalFungi variants of 3.6 could not be treated that way and were harmonized instead by removing the conserved 5.8S remnant (5^*′*^) and LSU remnant (3^*′*^) with `cutadapt` v4.4 [13], the identical command for both. The 5^*′*^ and 3^*′*^ adapters were specified independently rather than as a linked pair, with a 15% error tolerance and a minimum retained length of 50 bp. All three routes end at ITS2, which is what the identity comparisons require, but they are not the same operation and their effect varies with the starting state: `cutadapt` trimmed 98.3% of the soil ASVs and only 3.3% of the GlobalFungi variants, which that project already delimits to ITS2 by its own extraction. Residual disagreement over exactly where each pipeline places the boundary is absorbed by scoring coverage over the shorter sequence rather than requiring mutual full-length identity (§3.6).

### 3.2 Leave-one-genus-out benchmark construction

From the reference backbone we built two held-out splits. *Leave-one-genus-out* (LGO) withholds a query’s entire genus from the references, giving 4,444 novel-genus ITS2 queries that constitute the novel-positive class. *Leave-one-species-out* (LSO) withholds only the species while retaining its genus, giving 3,170 known-genus queries as the known-negative class. Split construction guarantees that each query’s true higher-rank lineage remains present among the references, so that family-level placement is in principle recoverable and novelty is genuinely a property of the genus rather than of the whole lineage. Splits were generated with a fixed random seed and are regenerated by the harness from public data.

### 3.3 Novelty scoring, separability and calibration

Each query was searched against the reference with `vsearch` v2.31.0 [19] (`usearch_global`, best hit, identity floor 0.5); the best-hit percentage identity is the novelty signal, higher identity implying a closer known relative. Separability of novel from known queries was quantified by the area under the ROC curve on the negated identity.

To convert a score into an error-controlled decision we used inductive conformal anomaly detection [24]. Known queries were partitioned into a calibration set (1,571) and a test set (1,572); each test query received a conformal *p*-value equal to the fraction of calibration knowns at least as novel as it, and was flagged novel when *p ≤ α*. Under exchangeability of the calibration and test scores this bounds the false-novelty rate on genuine knowns at *α*. Nothing in the procedure inspects the score’s construction, so identity, a bootstrap consensus or a neural output probability pass through the same machinery; where a score is discrete the bound is attained conservatively rather than exactly, because ties inflate the *p*-value.

Alongside the empirical false-flag rate we report, for every setting, the detection rate on novel-genus queries at the same *α*. The two quantities answer different questions, only the first is guaranteed, and reporting the first alone would leave the central practical caveat of the method unstated.

### 3.4 Cross-score verification

To establish that the guarantee holds beyond the score it was developed on, we recalibrated the flag on three further scores taken from the companion study [17]: the family-rank bootstrap confidence of SINTAX [8] as implemented in vsearch, and the family-rank output probability of two pretrained neural classifiers distributed with MycoAI [20]. Those scores were computed on 5,222 UNITE queries, one per genus, stratified by whether a query’s genus lies in the classifiers’ training label space, and were available for both full-length ITS and the ITS2 subregion of the identical records; the calibration set in each case was a random half of the seen-genus queries (*n* = 1,665). We additionally recalibrated alignment identity against references of 16,352, 56,327 and 113,455 sequences to test sensitivity to reference composition. In every case the quantities of interest are the empirical rate at which the flag fires on held-out known-genus queries, which should approximate the nominal *α*, and the rate at which it fires on novel-genus queries, which is not guaranteed by anything.

### 3.5 Outgroup control and application data

As a maximal-novelty control we scored 5,000 SSU rRNA sequences (SILVA 138.2 SSURef NR99) against the fungal ITS2 reference and passed the resulting identities through the flag. A correctly calibrated flag should assign the novel verdict to essentially all, SSU and ITS2 being non-homologous. We then applied the full pipeline to a soil-fungal dataset (Bio Project PR-JNA704912; 234 fungal ITS2 amplicon runs isolated from a dual-marker submission via the ENA `experiment_title field`). Reads were primer-trimmed with `cutadapt` and denoised with DADA2 [7] (`truncLen`= 0, to preserve ITS2 length variation), yielding 5,521 amplicon sequence variants (ASVs) that were ITS2-harmonized and scored as above.

### 3.6 Cross-study recurrence analysis

To assess whether flagged-novel ASVs recur in independent studies we searched the ITS2 sequence-variant library of GlobalFungi [23], as deposited in 2020 with that database’s description and spanning the 178 studies of that release, against all 5,521 ASVs, so that flagged and non-flagged ASVs share one search and one denominator. Two arms were run against that database, after identical ITS2 primer removal: the 50,266,003 variants carrying no assigned UNITE species hypothesis, and the 26,096 representatives of hypotheses that do carry a name. Both used `vsearch` with `–strand both`, an identity floor of 0.94, and `–maxaccepts 20 –maxrejects 0`; the last matters because with the default single accepted hit a global variant matching both a flagged and a non-flagged ASV is reported against only one of them, which would bias the comparison between the groups for purely mechanical reasons. Because the two datasets define the ITS2 window differently, recurrence was scored over the global variant rather than the ASV: an ASV recurs if *any* global variant aligns to it at the stated identity over at least 90% of that variant’s length, taken from vsearch’s qcov field. Any rather than best hit, since the question is whether the sequence occurs elsewhere and not whether its most similar global relative clears the threshold. The two arms are not symmetric and are not compared with each other: one searches tens of millions of individual variants and the other twenty-six thousand cluster representatives.

## 4 Results

### 4.1 Alignment identity separates novel from known only moderately

Novel-genus queries reached a mean best-hit identity of 77.96% against the ITS2 reference, against 88.37% for known-genus queries, a gap of roughly 10.4 percentage points yielding AU-ROC = 0.745. Identity therefore carries real but incomplete novelty signal, and no single identity threshold separates the two classes cleanly. This is precisely the situation a calibrated decision rule is meant to handle: the question is not where the classes divide, since they overlap, but how to act at a stated error rate given that they do.

### 4.2 The flag achieves calibrated error control

The empirical false-flag rate tracks the nominal *α* closely across two orders of magnitude (Table 1, Figure 1). At *α* = 0.05 the flag detects 19.2% of true novel-genus queries while holding false flags at 4.3%: specific but conservative, as an AUROC of 0.745 requires. The identity cutoffs the procedure derives are worth reading alongside the conventional 97%: to hold the false-novelty rate at 5% on this reference the implied cutoff is 64.8% identity, and the customary value would flag a large majority of known-genus queries. The gap should not be read as evidence that 97% is the wrong convention in general. It follows from what the calibration set contains. Leave-one-species-out withholds the query’s species, so a known-genus query’s best hit is a congener rather than a conspecific and the known-class identity distribution carries a correspondingly long left tail; a reference with dense species-level coverage of the queries at hand would calibrate to a substantially higher cutoff. That dependence is the practical content of calibration rather than an objection to it. The threshold achieving a stated rate is a joint property of the reference and the queries, is not knowable in advance, and does not travel between datasets even when the score does.

**Table 1:**
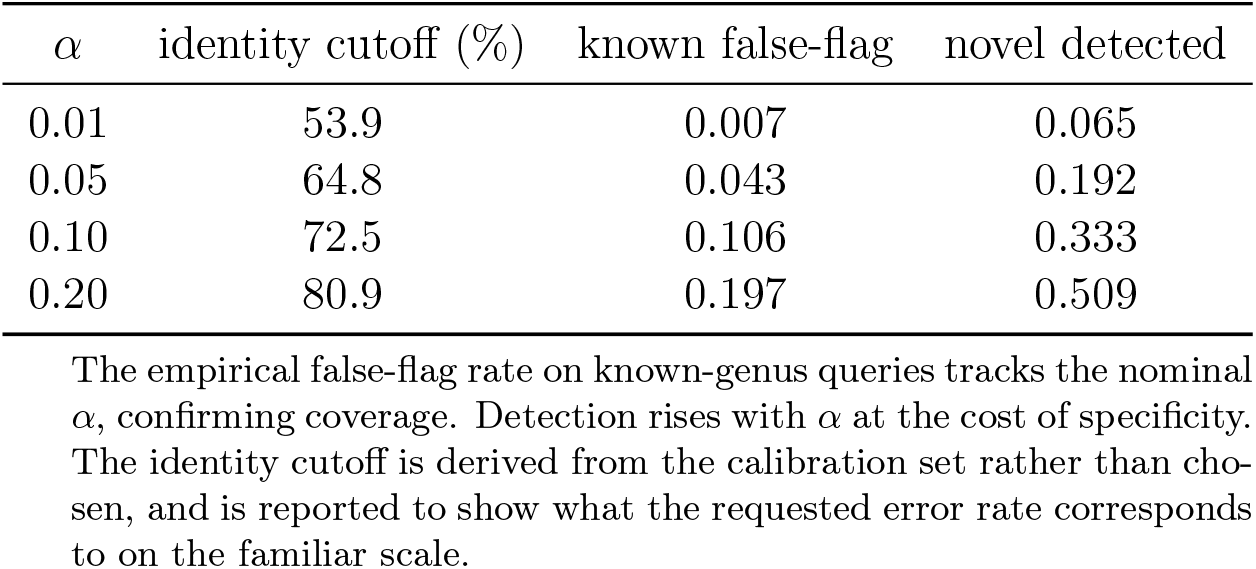
Conformal novelty flag across operating points (ITS2, alignment identity).

**Figure 1:**
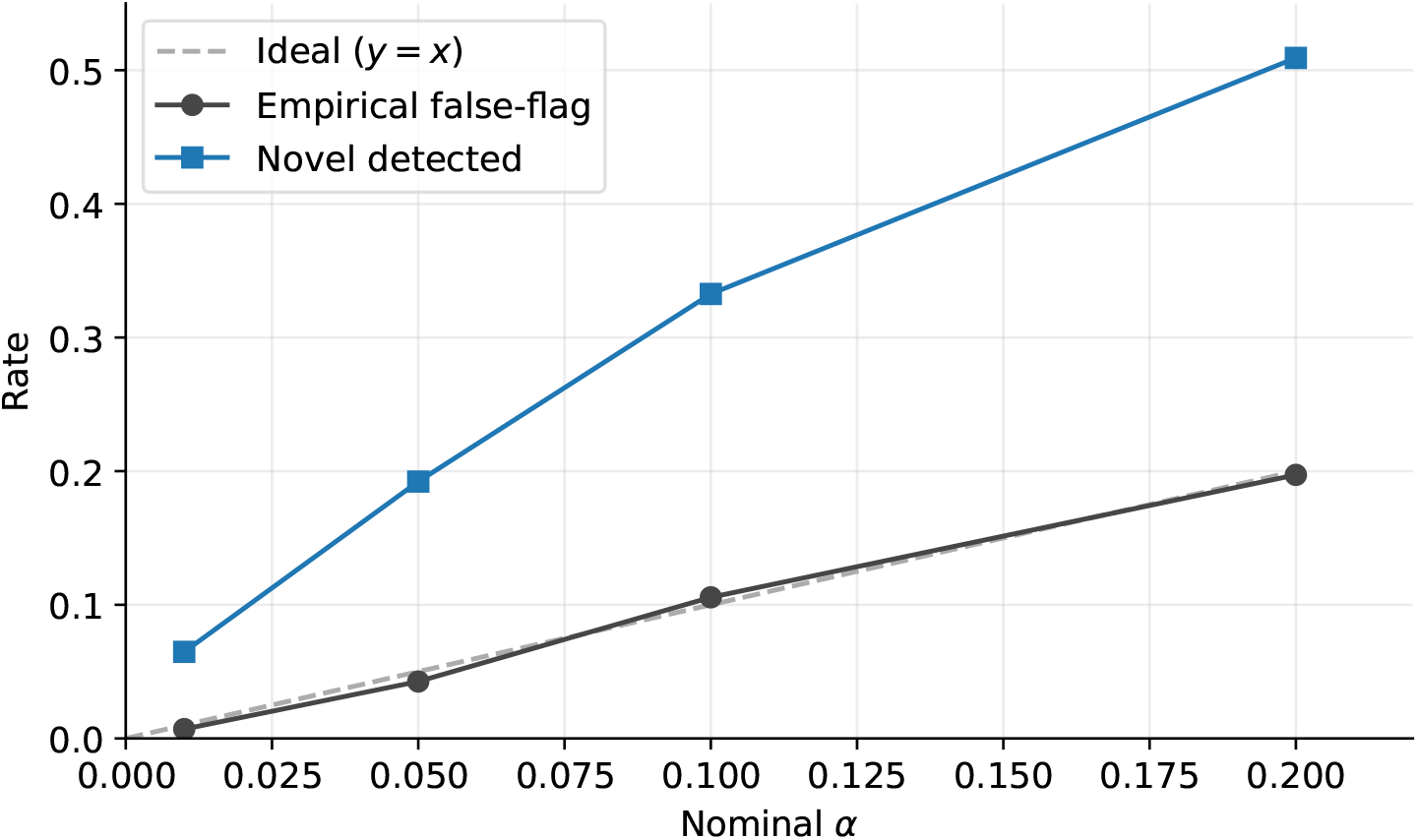
Conformal calibration on ITS2. The empirical false-novelty rate on known-genus queries tracks the nominal *α*, points falling on the identity line, confirming the distribution-free coverage guarantee. Detection of true novel-genus queries, the upper curve, is limited by the moderate separability of the underlying score, so the flag is specific but conservative, which is the appropriate posture for an abstention mechanism where a false novelty call is more costly than a missed one.

### 4.3 The flag behaves correctly across a known, novel and outgroup gradient

The outgroup anchors the extreme of the novelty scale (Figure 2): a completely non-homologous marker is recovered as 99.9% novel, confirming that the flag’s novel calls are not artefacts of the permissive search floor. Together the three populations trace the expected monotonic gradient from known (4.3%) through novel-genus (19.2%) to outgroup (99.9%), and the two mislabelled fungal sequences the flag declined to call novel are a reminder that its verdicts are interpretable at the level of individual sequences.

**Figure 2:**
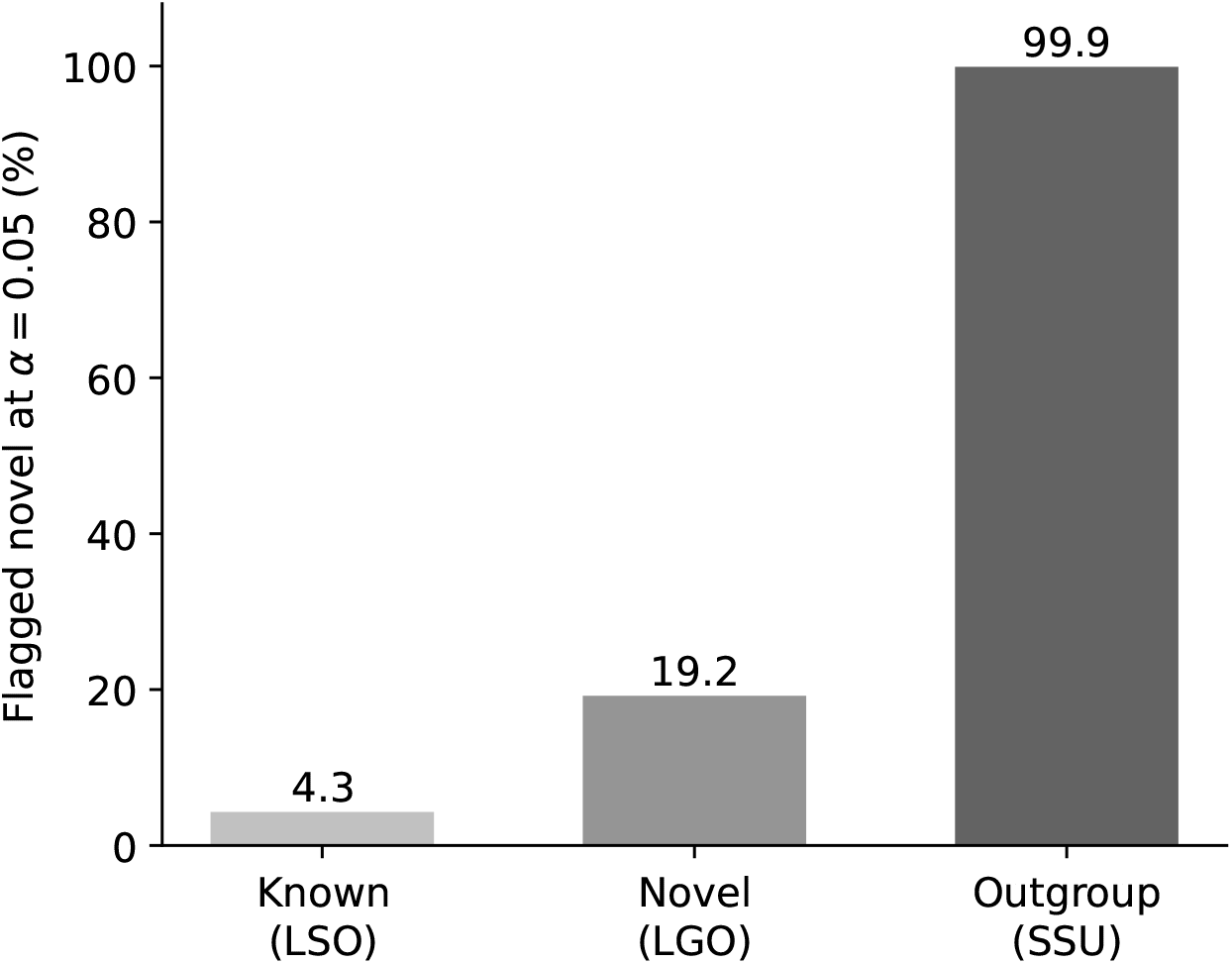
Novel-call rate at *α* = 0.05 across three populations spanning the novelty spectrum. The flag fires rarely on known-genus queries (4.3%, the calibrated false-flag rate), modestly on novel-genus queries (19.2%, its conservative power at this operating point), and on 99.9% of a non-fungal SSU outgroup. Of 5,000 outgroup sequences, 4,996 were flagged novel; the four exceptions were two chance alignments near the cutoff and two fungal sequences (Cordycipitaceae and *Trametes versicolor*) mislabelled as SSU in the source subset, identified on inspection.

### 4.4 Coverage transfers across scores; power does not

A guarantee that held only for the score it was developed on would be of limited use, since scoring methods differ and proliferate. We therefore recalibrated the flag on three further scores and across two amplicon regions and three reference depths (Table 2). All eleven settings produced empirical false-flag rates within 1.1 percentage points of the nominal *α* = 0.05, the extremes being 0.041 for a discrete bootstrap consensus and 0.061 for a neural softmax. The scores involved differ in construction, in scale and in what they measure; a threshold calibrated for one has no interpretation under another, whereas *α* retains its meaning throughout. Coverage also held across a sevenfold change in reference size, which matters because the reference is the component a practitioner is most likely to alter and the one a fixed identity threshold is most sensitive to.

**Table 2:** Empirical false-flag rate on held-out known-genus queries and detection rate on novel-genus queries, both at a nominal *α* = 0.05, for scores of differing provenance. Coverage is guaranteed and holds throughout; detection is not guaranteed and varies more than tenfold.

| score | setting | false-flag | detection | AUROC |
| --- | --- | --- | --- | --- |
| alignment identity | ITS2, LGO benchmark | 0.043 | 0.192 | 0.745 |
| alignment identity | full ITS, reference 16,352 | 0.050 | 0.260 | 0.836 |
|  | full ITS, reference 56,327 | 0.053 | 0.453 | 0.872 |
|  | full ITS, reference 113,455 | 0.046 | **0.533** | 0.887 |
|  | ITS2, reference 54,108 | 0.048 | 0.427 | 0.866 |
| SINTAX bootstrap | full ITS | 0.041 | 0.313 | 0.813 |
|  | ITS2 | 0.055 | 0.344 | 0.785 |
| classifier $P(\text{family})$, CNN | full ITS | 0.061 | 0.205 | 0.706 |
|  | ITS2 | 0.046 | **0.061** | 0.503 |
| classifier $P(\text{family})$, BERT | full ITS | 0.049 | 0.178 | 0.688 |
|  | ITS2 | 0.042 | **0.052** | 0.541 |

Scores other than alignment identity on the LGO benchmark, and the AUROC values, are taken from the companion study [17]. All eleven false-flag rates fall within 1.1 percentage points of nominal; detection ranges from 0.052 to 0.533. The two ITS2 classifier rows, in which detection barely exceeds the false-flag rate, are the cases where the flag is valid and uninformative simultaneously. Conservative coverage is expected where a score is discrete, as the SINTAX bootstrap is. Detection is comparable within each block but not between them, the first row being the leave-one-genus-out benchmark of this paper and the remainder the companion’s one-per-genus queries, and the gap follows entirely from the known class each calibrates on. Leave-one-species-out knowns have a median best-hit identity of 92.1% and a fifth percentile of 65.3%, the latter being the cutoff the procedure derives; the companion’s seen-genus queries retain conspecifics and reach a median of 99.5% with a fifth percentile of 79.6%. Applied to a novel class whose median identity is 82.2%, a cutoff near 80% catches most of the lower half and one near 65% catches little of it, which is the whole of the difference between 42.7% and 19.2%. This is the dependence set out in §4.2 measured rather than asserted, and it is why $\alpha$ travels between datasets and a cutoff does not.

Reporting detection alongside coverage is what makes the scope of the guarantee legible. Across the same eleven settings, under a false-flag rate that never departs from nominal by more than a percentage point, detection ranges from 0.052 to 0.533, a better than tenfold spread. The guarantee is on the error rate; nothing in it constrains how many genuine novelties are found. Detection instead follows the separability of the underlying score, though not exactly: it rises with reference depth in step with AUROC (0.260, 0.453, 0.533 against 0.836, 0.872, 0.887, all within the second block of the table), and the ordering of methods by detection broadly matches their ordering by AUROC, but there are inversions, since AUROC integrates the whole ROC curve whereas detection at a fixed *α* depends only on the left tail of the known-class score distribution relative to the novel class. Detection is therefore an empirical property of each deployment and must be measured rather than inferred from a summary statistic.

The two ITS2 classifier rows make the point concretely rather than hypothetically. The convolutional model’s *P* (family) separates novel from known genera at AUROC 0.503 on ITS2, which is chance; the flag calibrated on it holds a false-flag rate of 0.046, within half a point of nominal, and detects 6.1% of true novelties, barely above the rate at which it fires on knowns. The transformer behaves the same way, 0.042 against 0.052. In both cases coverage is essentially exact and the resulting decision is worthless: the flag fires at approximately *α* on both classes, which is what a valid test built on an uninformative statistic must do. Coverage and power are therefore independent properties, only the first is guaranteed, and Table 2 should be read as a validity check across scores rather than as a recommendation to wrap whichever score happens to be available. The practical counsel is to calibrate whatever score you have, and then to measure its detection rate before relying on it.

### 4.5 Application: dark matter in soil fungal communities

Applied to real soil data the flag identifies 20.7% of ASVs as novel at *α* = 0.05 (Figure 3), a dark-matter fraction in line with independent soil-fungal surveys, and one that now carries a stated error rate rather than resting on a chosen cutoff. As a determinism check the 234 samples were randomly split into two halves and denoised independently; per-sequence verdicts agreed perfectly (Cohen’s *κ* = 1.0), confirming that the flag is a deterministic function of sequence and reference and introduces no stochastic instability.

**Figure 3:**
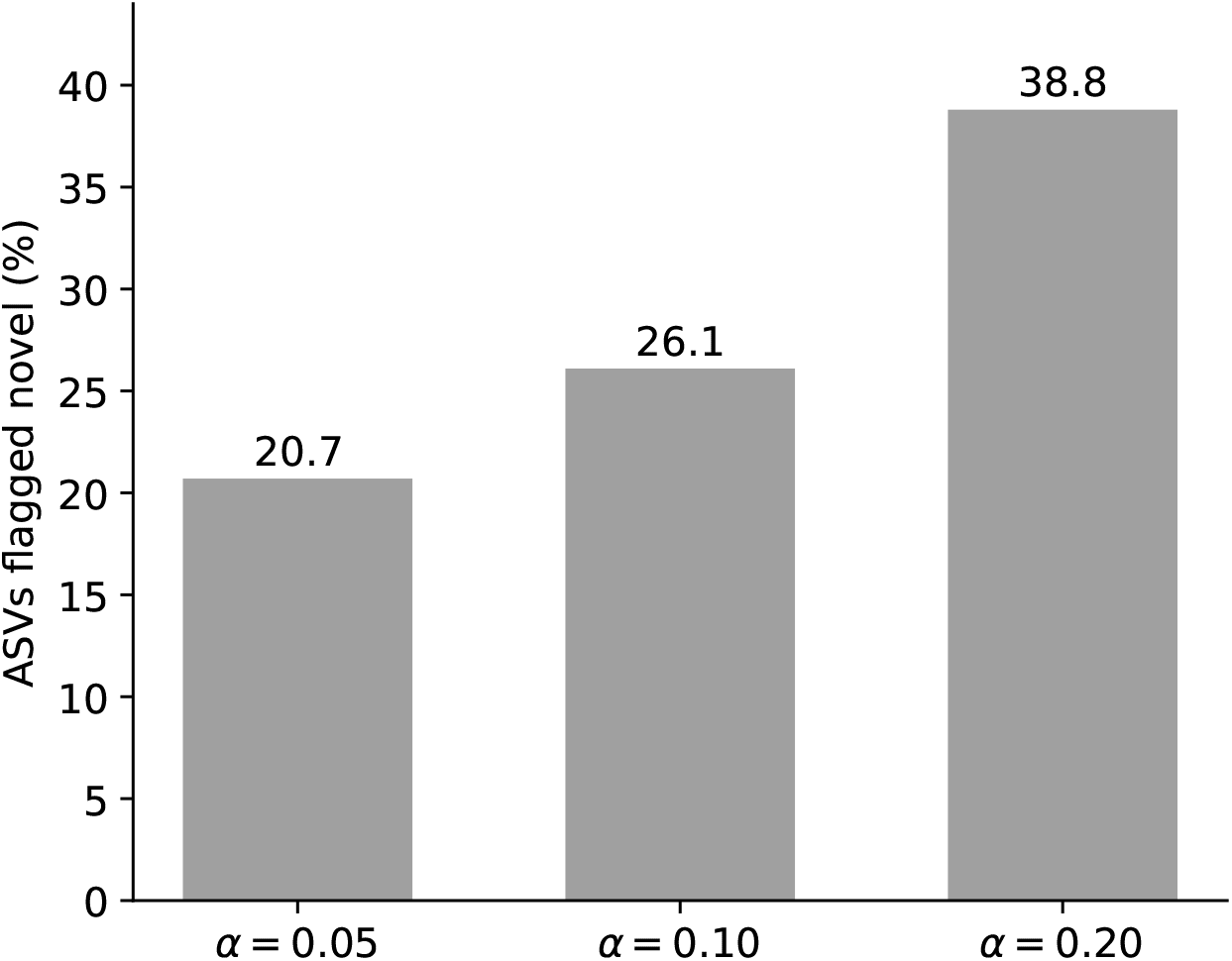
Novel fraction of 5,521 soil fungal ASVs across operating points. At *α* = 0.05, 20.7% of ASVs are flagged novel, rising to 26.1% at *α* = 0.10 and 38.8% at *α* = 0.20, consistent with prior soil-fungal dark-matter estimates. Of the 5,521 ASVs, 5,075 obtained a reference hit above the identity floor and 446 did not, the latter being novel by construction.

Determinism is not freedom from error, and denoising was run without an abundance or prevalence filter, so an error-derived variant with depressed identity against the reference is flagged novel by construction. Of the 5,521 ASVs, 3,529 occur in a single sample, and novel calls are enriched among them: restricting to the 1,992 present in at least two samples lowers the novel fraction from 20.70% to 16.27% at *α* = 0.05, from 26.08% to 20.28% at *α* = 0.10 and from 38.76% to 33.99% at *α* = 0.20; a ≥ 10 read filter gives 16.38% and both together 15.94%. We report both bounds rather than choosing between them, because neither is the quantity of interest on its own. The unfiltered fraction includes residual denoising error and the filtered one discards genuinely rare taxa, and §4.6 shows the discarded set is not simply error: 48.4% of the single-sample ASVs flagged novel recur near-identically in independent studies worldwide, which an error-derived variant does not do.

### 4.6 Flagged sequences recur globally but are rarely named

To test whether flagged-novel sequences are genuinely rare or merely absent from the curated reference, we asked whether they recur in the ITS2 variant library of Global Fungi [23], a compilation spanning the 178 studies of the release searched here and considerably more in subsequent ones, searching both its unnamed and its named partitions against all 5,521 ASVs (3.6) so that flagged and non-flagged ASVs are scored on one search (Table 3).

**Table 3:**
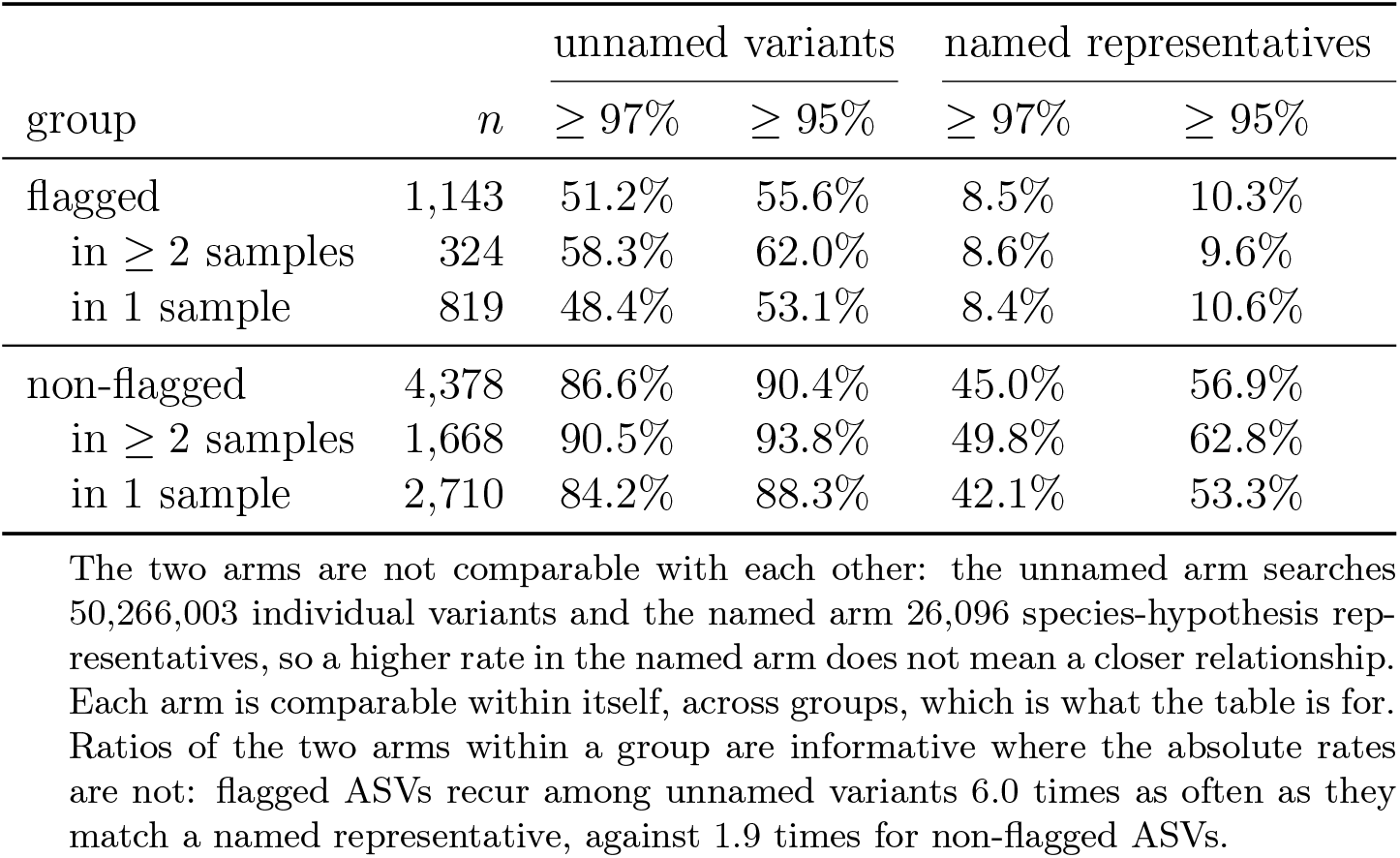
Cross-study recurrence of soil ASVs in the GlobalFungi ITS2 variant library. Percentage of ASVs in each group matching at least one global variant at the stated identity over at least 90% of that variant.

| group | *n* | unnamed variants |  | named representatives |  |
| --- | --- | --- | --- | --- | --- |
| | | $\geq 97\%$ | $\geq 95\%$ | $\geq 97\%$ | $\geq 95\%$ |
| flagged | 1,143 | 51.2% | 55.6% | 8.5% | 10.3% |
| in $\geq 2$ samples | 324 | 58.3% | 62.0% | 8.6% | 9.6% |
| in 1 sample | 819 | 48.4% | 53.1% | 8.4% | 10.6% |
| non-flagged | 4,378 | 86.6% | 90.4% | 45.0% | 56.9% |
| in $\geq 2$ samples | 1,668 | 90.5% | 93.8% | 49.8% | 62.8% |
| in 1 sample | 2,710 | 84.2% | 88.3% | 42.1% | 53.3% |

The two arms are not comparable with each other: the unnamed arm searches 50,266,003 individual variants and the named arm 26,096 species-hypothesis representatives, so a higher rate in the named arm does not mean a closer relationship. Each arm is comparable within itself, across groups, which is what the table is for. Ratios of the two arms within a group are informative where the absolute rates are not: flagged ASVs recur among unnamed variants 6.0 times as often as they match a named representative, against 1.9 times for non-flagged ASVs.

Of the 1,143 flagged ASVs, 585 (51.2%) matched at least one unnamed global variant at 97%≥ identity over at least 90% of that variant, and 636 (55.6%) at ≥ 95%; the corresponding figures for the 4,378 non-flagged ASVs are 86.6% and 90.4%. The named arm separates the two groups sharply and in the opposite direction: 8.5% of flagged ASVs match a named species-hypothesis representative against 45.0% of non-flagged ones, and 10.3% against 56.9% at the relaxed threshold. That separation is close to definitional, flagged ASVs being those with low identity to UNITE and the representatives being UNITE-derived, so we read it as a check that the flag does what it claims rather than as a finding about the sequences.

Abundance does not explain the flagged recurrence. Flagged ASVs present in at least two samples recur among unnamed variants at 58.3% against 48.4% for those seen in a single sample, a gap of 9.9 percentage points against 6.3 for the same contrast among non-flagged ASVs. In the named arm the prevalence effect appears only among non-flagged ASVs, 49.8% against 42.1%, and is flat among flagged ones, 8.6% against 8.4%, which is what one expects if flagged sequences are absent from the reference rather than merely uncommon. Global recurrence on its own does not distinguish flagged from known sequences, and it would have been convenient but wrong to report that it does. Non-flagged ASVs recur among unnamed variants far more often than flagged ones, 86.6% against 51.2%, which has a mundane explanation: the unnamed partition holds fifty million sequences drawn from real soil communities, and an abundant, widely distributed taxon has more opportunities to appear among them than a rare one. The prevalence effect within each group points the same way.

What the two arms establish jointly is narrower and, we think, the claim worth making. Half of the sequences the flag declares novel recur near-identically in independent studies worldwide, while fewer than one in ten matches any named species hypothesis: a six fold skew towards the uncatalogued partition, against a 1.9-fold skew for the sequences the flag passes. They are present in soils across the world and named in almost none of them. That is difficult to reconcile with their being local curiosities or denoising artefacts, and straightforward to reconcile with their being real fungal lineages that reference databases have not yet described.

The single-sample comparison closes the artefact objection directly. Of the flagged ASVs detected in exactly one of the 234 samples, 48.4% recur near-identically in independent studies on other continents, which a sequencing or denoising error cannot do. The fraction that an abundance filter sets aside in 4.5 is therefore a mixture of error and genuinely rare taxa rather than error alone, and the filtered 16.27% is a floor on the dark-matter fraction rather than a correction to it.

The ASVs that do recur are the defensible shortlist rather than the whole flagged set: they are demonstrably widespread, demonstrably unnamed, and selected by a procedure with a stated error rate, which makes them natural priority targets for formal characterization. The remainder are genuinely restricted, differ enough in the captured ITS2 window to escape detection here, or are error-derived variants that no independent study will ever recover; distinguishing the three is beyond what the present data can resolve.

## 5 Scope and limitations

Several limitations bound these results. First, novelty is defined relative to a chosen reference: a sequence flagged novel against UNITE may match sequences in larger environmental databases, and indeed half do. The reported fractions measure novelty with respect to the curated reference, which is the quantity a practitioner using that reference needs, but it is not a claim about absolute novelty. Second, the guarantee concerns the false-novelty rate alone. Detection varied more than tenfold across the settings in Table 2 under an essentially constant error rate, so a calibrated flag is not by itself evidence of a useful one and its detection rate must be measured for each deployment. Third, the moderate separability of alignment identity (AUROC 0.745) is a genuine ceiling of the score rather than a defect of calibration; raising detection requires a more informative representation, not a different threshold, and the flag is built to accept one. Fourth, the coverage guarantee rests on exchangeability between calibration and test scores. Our splits satisfy this by construction, but a practitioner calibrating on one dataset and applying the flag to another with a different length distribution or sequencing protocol may not, and the empirical rate should be re-checked in that situation. Fifth, the cross-study recurrence analysis compares datasets with different ITS2 window definitions and different UNITE versions, so it is scored over the shorter variant’s extent; this is conservative for detecting recurrence but precludes a strict full-sequence identity claim. Sixth, the flag detects novelty rather than error: it fires on novel genera whether or not a downstream assignment happened to be correct, which is the appropriate behaviour for surfacing uncatalogued diversity but not for auditing a classifier’s mistakes. Seventh, denoising was run without length truncation to preserve ITS2 length variation, and residual error-derived variants would be scored as novel by construction, so the application fraction is reported both unfiltered and restricted to ASVs present in at least two samples, 20.70% and 16.27% at *α* = 0.05, which bracket rather than resolve the residual-error question (§4.5). Finally, the UNITE species-hypothesis namespace [11] differs between releases and derived products, so cross-database comparison was performed at the sequence level rather than by shared identifiers.

## 6 Benchmark details and reproducibility

Analyses used vsearch v2.31.0, `cutadapt` v4.4 and DADA2 v1.38.0 on R 4.5.3. The harness is organised in five stages:

~~~
calibrate      prep_split.py, score_recovery.py, conformal_novelty.py soil data dada2_its2.R, dada2_half.R, score_asvs_conformal.py, asv_abundance.R, make_flagged_fasta.py
cross-study   gf_recurrence.sh, frey_sh_verdicts.py, gf_prevalence.py
selection     pick_studies.py
figures       make_figures.py
~~~

The deposit contains these and no others. `dada2_half.R` is the split-half determinism check and `make_flagged_fasta.py` regenerates the flagged set from the p-value column, so that it cannot drift from the verdict table. `frey_sh_verdicts.py` and `gf_prevalence.py` compare cross-study prevalence at the level of UNITE species hypotheses rather than of sequences. That comparison is not reported here, only 124 of the flagged variants carrying a species hypothesis to look up, and it is the only part of the harness reading a different GlobalFungi release; both are deposited for completeness and no result in this manuscript depends on them. `gf_recurrence.sh` replaces an analysis that previously had no script, which is why earlier recurrence figures could not be reproduced from the deposited hits. Its summary mode re-derives the recurrence table from that output without repeating the search, and because the search floor is 0.94 any identity threshold above it is a post-hoc filter rather than a search parameter, so the reported rates can be recomputed, or computed at a further threshold, from the hits alone:

~~~
ARM=summary bash gf_recurrence.sh
~~~

The reference was the UNITE general release `sh_general_release_dynamic_19.02.2025`; the calibration reference set comprised 33,982 sequences drawn from the 99,808-sequence ITS2 pool by leave-one-species-out. An ASV is flagged novel when its conformal p-value is at most *α*; the identity cutoff the procedure implies is reported for interpretation only and is used as a threshold nowhere in the analysis. Representative commands:

~~~
prep_split.py --input unite.ITS2.fasta --out out_lso_its2 --mode lso vsearch --usearch_global queries.fasta --db refs.fasta --id 0.5 \
  --maxaccepts 1 --top_hits_only --userfields query+target+id
conformal_novelty.py --known lso.hits --novel lgo.hits \
  --alphas 0.01,0.05,0.10,0.20
~~~

The soil dataset (BioProject PRJNA704912) was streamed from the European Nucleotide Archive and the GlobalFungi variant library from the figshare collection doi:10.6084/m9.figshare.c.4915392. Scores for the cross-score verification were produced by the companion study’s harness [17] and are deposited alongside it. All inputs, commands and analysis scripts are included in the repository so that every table and figure here can be regenerated from public data.

## 7 Discussion

The contribution of this paper is narrow and, we think, useful: it replaces a chosen cutoff with a chosen error rate. Novelty detection in fungal metabarcoding has long proceeded by thresholding a score at a conventional value, and that practice works better than it might: the conventions in use are not erratic. But it cannot answer the question a practitioner actually has, which is what rate of mistakes a given decision entails, and it offers no way to move to a different rate deliberately. The conformal flag answers both, and the identity cutoffs it derives make the point vividly. To hold the false-novelty rate at 5% against our leave-one-species-out calibration set the implied cutoff is 64.8% identity, nowhere near the customary 97%. The distance between those two numbers is not an argument that the convention is wrong; it is a measure of how strongly the correct cutoff depends on the species-level completeness of the reference at hand, and therefore of how little a threshold carries its error rate with it when it moves between datasets.

That the guarantee transfers is what makes it practical rather than merely principled. Across eleven settings spanning four scores of quite different construction, two amplicon regions and a sevenfold change in reference size, the empirical rate stayed within about one percentage point of nominal. This matters because the field’s scoring landscape is changing: alignment identity, bootstrap consensus and neural probabilities are all in current use, and a wrapper that treats them identically outlives any particular one of them.

It is equally worth being clear about what calibration does not do, and our own results supply the illustration. Under a false-flag rate that never departed from nominal by more than a percentage point, detection across those same eleven settings ranged from 5.2% to 53.3%. Two of them, both neural classifiers applied to ITS2, detected genuine novelties at almost exactly the rate at which they fired on knowns, which is what a valid test built on a statistic carrying no signal must do. Coverage bounds the cost of a decision; it does not make the decision informative. Any deployment of this flag should therefore report its detection rate alongside its error rate.

The biological result gives the flagged material weight, and it now rests on a contrast rather than a bare proportion. Sequences the flag declares novel match a named Global Fungi species hypothesis 8.5% of the time against 45.0% for the sequences it passes, while recurring among the library’s unnamed variants at 51.2%: a sixfold skew towards the uncatalogued partition, where the sequences it passes show a 1.9-fold skew. They are present in soils worldwide and named in almost none of them. That is difficult to reconcile with their being artefacts or local curiosities and straightforward to reconcile with their being real fungal lineages reference databases have not yet described. The abundance stratification closes the obvious objection: 48.4% of flagged ASVs seen in exactly one sample recur near-identically in independent studies on other continents, which a denoising error cannot do, so the single-sample fraction is a mixture of error and genuinely rare taxa rather than error alone.

The flag therefore does more than tell an analysis when to abstain. It produces a shortlist: sequences that are demonstrably widespread, demonstrably unnamed, and selected at a stated error rate. For a field whose central obstacle is that most of what it recovers cannot be named, converting dark matter from a residue into a prioritized set of description targets seems to us the more consequential use. What such a shortlist can then support has been a contested question [12, 14], but the ground has moved: thirty fungal lineages have now been formally described from environmental samples and DNA, with the terms *nucleotype* and *legitype* proposed for DNA-derived types [21]. A calibrated shortlist is on that reading not only a research priority list but a candidate input to description itself. Our claim remains the narrower one, that these are the sequences a description effort should start from, on whatever nomenclatural basis the field settles.

Where the flag is currently limited is in power rather than in validity. At *α* = 0.05 on the leave-one-genus-out benchmark it detects under a fifth of genuinely novel genera, a ceiling imposed by the separability of the underlying score and not by the calibration machinery, and the cross-score table shows how much room that leaves: the same procedure reaches 53.3% detection at the same error rate when given a more separable score. Because the procedure is score-agnostic, the natural route to improvement is a better novelty score rather than a different decision rule. A representation trained with held-out genera in its objective, so that unseen genera fall near their true relatives, would raise detection at the same guaranteed error rate, and could be benchmarked on the same harness against the 0.745 baseline reported here.

## Author contributions

Aaron O’Brien conceived the study, implemented the harness and the conformal calibration, ran all analyses and wrote the original draft. César Marín critically reviewed and revised the manuscript, contributing edits throughout and additions to the framing of fungal dark taxa and to the literature cited. Pilar Parada acquired the funding (CORFO 23PTECCC-247149) and supervised the project. All authors read and approved the submitted version.

## Data and code availability

Benchmark splits, the ITS2 harmonization pipeline, the conformal calibration code and the per-query verdict tables every value here is computed from are available under the MIT license at https://github.com/ayobi/its-novelty. The GlobalFungi search output is not deposited, being 1.3 GB and regenerable from public data by the deposited script. Soil data: NCBI BioProject PRJNA704912. Reference: UNITE general FASTA release for Fungi, version 19.02.2025, doi:10.15156/BIO/3301229. Outgroup: SILVA 138.2 SSURef NR99. GlobalFungi variant library: Baldrian et al., figshare collection, doi:10.6084/m9.figshare.c.4915392, file `GlobalFungi_ITS_variants.zip` as deposited on 5 April 2020 alongside the database descrition, containing 131,446,674 ITS variants. Every GlobalFungi result reported here derives from that file; the release-5 abundance tables distributed separately are read only by the two scripts whose output is not reported (§6).

## Acknowledgements

This work was supported by CORFO grant 23PTECCC-247149 to Pilar Parada. César Marín thanks Fondecyt Regular Project No. 1240186 (ANID, Chile). We thank the Centro de Biotecnología de Sistemas for compute resources.

## Use of generative AI

Anthropic’s Claude and GitHub Copilot were used in preparing this work: to draft and revise manuscript text, to write and debug the analysis and figure scripts deposited with it, and to check bibliographic details. All analyses were run by the authors, all reported values were verified against the primary output, and the authors are responsible for the content of the manuscript.

## Notes

https://github.com/ayobi/its-novelty

### Competing Interest Statement

The authors have declared no competing interest.

### Summary of Updates

Revision and edits from Cesar, as well as fixes to typos etc

